# A spatially explicit and mechanistic model for exploring coral reef dynamics

**DOI:** 10.1101/2020.04.11.036913

**Authors:** Bruno S. Carturan, Jean-Philippe Maréchal, Corey J. A. Bradshaw, Jason Pither, Lael Parrott

## Abstract

The complexity of coral-reef ecosystems makes it challenging to predict their dynamics and resilience under future disturbance regimes. Models for coral-reef dynamics do not adequately accounts for the high functional diversity exhibited by corals. Models that are ecologically and mechanistically detailed are therefore required to simulate the ecological processes driving coral reef dynamics. Here we describe a novel model that includes processes at different spatial scales, and the contribution of species’ functional diversity to benthic-community dynamics. We calibrated and validated the model to reproduce observed dynamics using empirical data from Caribbean reefs. The model exhibits realistic community dynamics, and individual population dynamics are ecologically plausible. A global sensitivity analysis revealed that the number of larvae produced locally, and interaction-induced reductions in growth rate are the parameters with the largest influence on community dynamics. The model provides a platform for virtual experiments to explore diversity-functioning relationships in coral reefs.

## 1. Introduction

Corals are the foundation species of coral-reef ecosystems and provide essential ecological functions for ecosystem resilience, as well as and services to millions of people worldwide (Moberg and Folke, 1999; Woodhead et al., 2019). The loss of coral cover and the change in species composition occurring in communities around the globe (Hughes et al., 2018; Torda et al., 2018) therefore alter the functioning of the entire ecosystem. For instance, the replacement of morphologically complex, but highly sensitive species, by simpler and more resilient species reduces the overall architectural complexity of reef habitats (Alvarez-Filip et al., 2009) — this simplification reduces the diversity and abundance of fish (Darling et al., 2017; Newman et al., 2015) and macroinvertebrates (Nelson et al., 2016) and the functions they provide (Pratchett et al., 2018). The change from species exhibiting fast rates of calcification to slower species reduces overall carbonate production (i.e., reef growth) (Alvarez-Filip et al., 2013; Perry et al., 2013) and disrupts bioerosion processes (Perry et al., 2014). The loss of reef complexity also reduces the dissipation of wave energy (Harris et al., 2018), increasing shoreline erosion (Graham and Nash, 2013; Sheppard et al., 2005) and the functioning of neighboring ecosystems by altering trophic links, energy and nutrient fluxes (Gillis et al., 2014; Harborne et al., 2006).

Disentangling the identity effect (effects of individual species on processes) from the diversity effect (shared effects of a collection of species on processes) is necessary to define which species, functional groups, or aspects of diversity are essential for maintaining ecological processes (Bellwood et al., 2019; Brandl et al., 2019). While the effect of such essential species on ecosystem function and the consequences of their loss has been widely reported (e.g., Hughes 1994, Alvarez-Filip et al 2009, Bellwood et al 2009), the relationship between species richness or functional diversity and ecosystem functioning is still poorly understood. Diversity is hypothesized to enhance functioning because of niche complementarity and facilitation (Bulleri et al., 2016; Loreau, 2000), but tests of this hypothesis with corals are scarce (but see McWilliam et al. 2018a).

Even less understood is the effect of diversity on the temporal variability (i.e., stability) and resilience (i.e., resistance and recovery) of community-level aggregate properties such as biomass, percentage cover, and calcification rate (Griffin et al., 2009). Higher diversity is hypothesized to promote ecosystem stability and resilience via asynchronous, independent population dynamics and compensatory population dynamics resulting from functional redundancy and response diversity — the latter is the insurance hypothesis (reviewed in Griffin et al. 2009 and McCann 2000). Tests of the insurance hypothesis are rare for coral reefs (but see Mellin et al. 2014 and Nash et al. 2016 for fish, and Zhang et al. 2014 and Clements and Hay 2019 for coral diversity).

Predicting how the reassembly of coral species in reefs alters ecosystem functioning is therefore a research priority to inform conservation management (Bellwood et al., 2019; Graham et al., 2014). To do this requires identifying the main ecosystem processes and associated functional traits affecting coral dynamics, and quantifying the relationship between the two (Bellwood et al., 2019; Carturan et al., 2018). Both virtual and physical experiments are also required (Brandl et al., 2019; Mcleod et al., 2019), where different aspects of diversity such as species and functional richness, and external factors such as disturbance regimes, larval connectivity, and grazing, are varied independently from one another to determine how they influence these dynamics. Unfortunately, physical experiments with coral species are unwieldy because of their slow growth rates and challenging environment in which they exist. However, models use simulation to overcome these challenges. To simulate realistically the effect of species and functional richness on ecosystem functioning, models should include: (*i*) links between functional traits and processes; (*ii*) biotic interactions such as spatial competition, and (*iii*) population dynamics representing demographic structure. Although some coral models have been developed (reviewed in Kubicek and Borell 2011, Weijerman et al. 2015), most implement only one or two of these aspects at most, and often with limited ecological detail. Importantly, no models have been developed to represent the breadth of species richness and functional diversity found in coral reefs in different regions of the world.

We present a new, spatially explicit, agent-based model representing benthic communities in tropical reefs composed of coral species and six functional groups of algae. Individual colonies grow, reproduce, compete for space, and respond to disturbances as a function of their size and trait-processes relationships, which we defined using eleven functional traits informed by published empirical data. The number of coral species present in the community can be varied without impacting model complexity and processing time. Functional diversity can be varied by sampling species from a set of 798 functionally realistic species, which we obtained by imputing missing trait data based on values for real species (Madin et al. 2016a). Our aim is to provide the full description of our model’s design, concepts, and capabilities. In the main text we present a streamlined description, and use the Appendices for details regarding: (1) traits and imputation of missing data (Appendix S1), (2) parameterization and processes implementation (Appendices S2 and S4), (3) calibration with empirical data (Appendix S3), (4) global sensitivity analysis (Appendix S6), and (5) hierarchically structured validation (Appendix S5).

## 2. Methods

### 2.1. Sources and software

We collected coral-trait data from coraltraits.org (Madin et al., 2016a) and other resources from the peer-reviewed literature (Appendix S1). We systematically verified and corrected coral-species nomenclature using the World Register of Marine Species as a reference. We used R (version 3.5.0, R-Core Team, 2017) to manipulate datasets, for statistical analyses, and to manage model simulations. We developed the model with the open-source, Java object-oriented programming language *Repast Simphony* 2.5.0 (North et al., 2013). We launched simulations using the R package `rrepast` 0.7.0 (García and Rodríguez-Patón, 2016) and `rJava` 0.9-10 (Urbanek, 2018). We used the R package `missForest` 1.4 (Stekhoven and Bühlmann, 2012) to impute missing trait data. We included phylogenetic information as a predictor using Huang and Roy’s (2015) phylogenetic supertrees to improve predictions; we manipulated the supertrees using the R packages `ape` 5.0 (Paradis and Schliep, 2019) and `phytools` 0.5-38 (Revell, 2012) (Appendix S1). We defined coral bleaching probabilities using the R packages `MuMIn` 1.40.0 (Bartón, 2017), `betareg` 3.1-0 (Cribari-Neto and Zeileis, 2010), `lme4` 1.1-15 (Bates et al., 2015), `lmtest` 0.9-35 (Zeileis and Hothorn, 2002) and `MuMIn` 1.40.0 (Bartón, 2017) (Appendix S4). For the model sensitivity analysis, we drew a Latin hypercube sample from the parameter space using the *randomLHS* function from the R package `lhs` 0.16 (Carnell, 2018), and we measured the influence of parameters on different response variables by fitting boosted regression trees using the *gbm.step* function from the R package `dismo` 1.1-4 (Hijmans et al., 2017).

### 2.2. Model description

We provide here a brief description of the model following the overview, design concepts, and details protocol (Grimm et al., 2010, 2006). A complete description of the parameterization and processes implementation along with a review of the supporting literature is available in Appendices S2 and S4.

#### 2.2.1. Purpose

The purpose of the model is to predict coral population dynamics as a function of cyclone and bleaching intensity and frequency, grazing pressure, larval connectivity, interspecific competitive interactions, and benthic community diversity (species richness and functional diversity).

#### 2.2.2. Entities, state variables, and scales

The model consists of grid-cell agents each representing 1 cm^2^, so that the benthic community is represented at a scale of organization smaller than the colony (equivalent to that of a polyp, although polyp size varies among species by several orders of magnitude) (Figure 1). Each agent is characterized by 32 variables that describe where the agent is in space (its position is fixed), its species identity and related functional characteristics, its age, the colony’s planar area and identification number, if it is bleached or was grazed recently, *et cetera*. Coral colonies and patches of algae are composed of multiple agents sharing the same variable values (except for their spatial coordinates). The state (variable values) of the agents changes as a function of the events happening in the reef. For instance, dislodgement is simulated by converting all the agents forming the dislodged colony into barren ground; a turf algae overgrowing a colony is simulated by converting the coral agents constituting the overgrown part into turf, but conserving the information about the colony (i.e., identification number, size, species, growth form).

**Figure 1.**
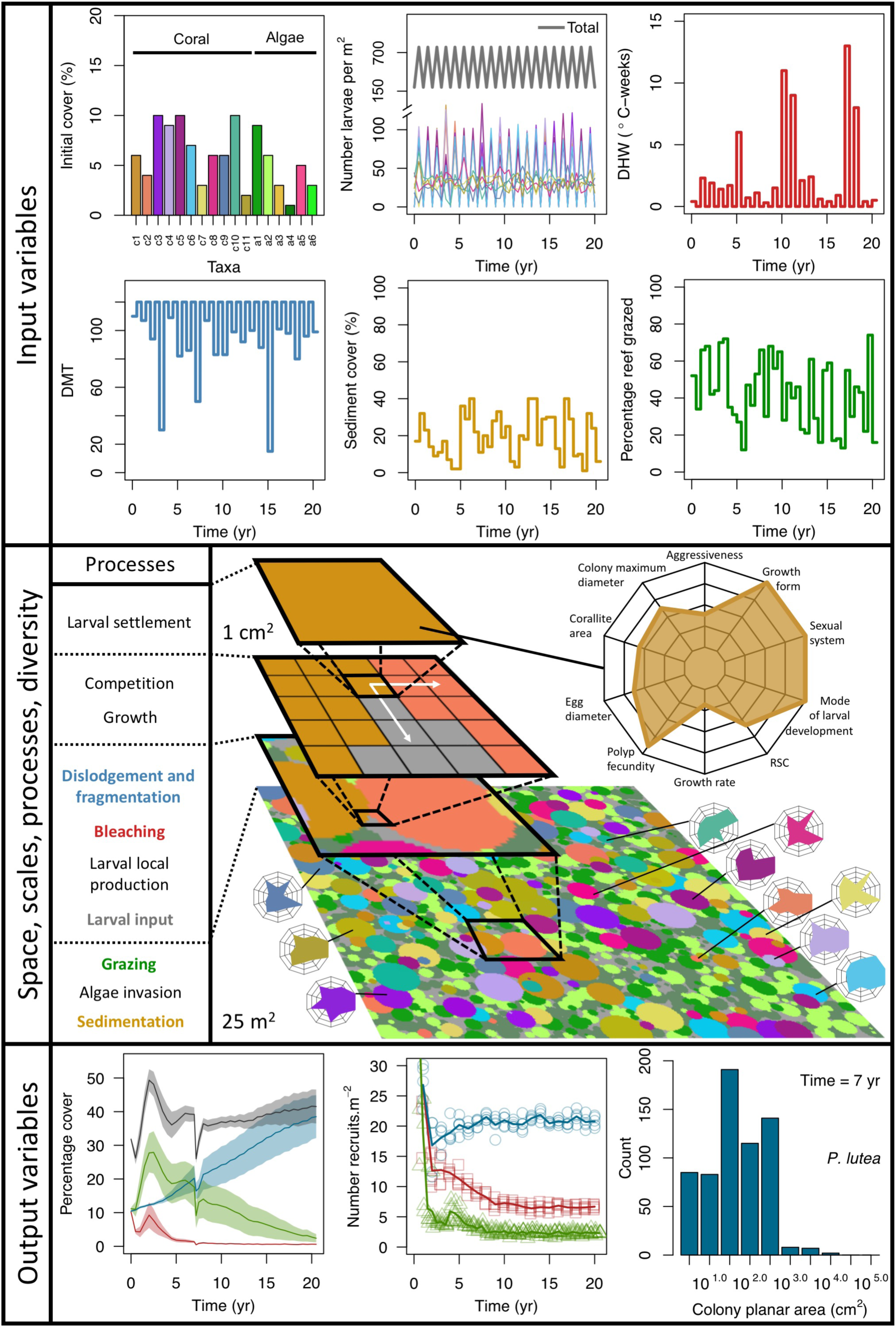
Description of the agent-based model. Six different variables as model inputs determine (*i*) initial community composition, (*ii*) number of larvae coming from the regional pool (total number divided among different species, with annual supply for spawning species, and biannual supply for brooding species), (*iii*) thermal stress in degree-heating weeks, (*iv*) hydrodynamic regime intensity expressed as dislodgment mechanical threshold (unitless), (*v*) sedimentation, and (*vi*) the percentage of reef grazed. All variables are inputs at every time period except for the initial community composition that is determined during initialization. The model represents a 25-m^2^ coral reef community and is composed of 1-cm^2^ cell agents. Once every time step, living agents (algae and corals) grow by converting their neighbouring agents within a certain radius (white arrows in middle panel). Different processes affect the community at different spatial scales. For instance, the grazing process lasts until the imposed percentage cover grazed over the entire reef is reached. In contrast, coral colonies are individually considered for dislodgment during hydrodynamic disturbance and a single agent is potentially converted into a new coral recruit when larvae successfully settle. Radar charts represent the functional characteristics of coral species (defined by a specific colour): each vertex corresponds to a functional trait and the coloured polygon indicates the trait values of the species (higher values are farther away from the centre of the web). At the end of each time step, the model provides for every taxon the percentage cover, the number of coral recruits, and the size of each colony (bottom panel).

The size of the reef and the length of a time step are changeable. We defined a 25 m^2^ reef for our simulations (i.e., 250,000 agents), which is usually the scale at which benthic communities are assessed in detail (e.g., Holbrook et al. 2018, Torda et al. 2018). We defined a 6-month time step because the empirical data we used to calibrate the model were collected biannually. Alternatively, it is possible to set three and 12-month time steps.

#### 2.2.3. Process overview and scheduling

Each time step is constituted of the following consecutive processes: (*i*) grazing, (*ii*) coral reproduction —locally and regionally produced larvae attempt to settle, (*iii*) thermal disturbance, which, if triggered, eventually causes colonies to bleach and/or die; (*iv*) dislodgement and fragmentation — the effect of waves and cyclones on certain colonies, (*v*) growth — each living agent, selected in a random order, attempts to convert its neighbouring agents within a certain radius to its own state, (*vi*) sedimentation — barren ground agents are converted to sand and *vice versa* until the desired sand cover is reached, (*vii*) algae invasion — the remaining ungrazed, barren-ground agents are converted into algae agents (Figure 2). The order at which processes *iii, iv*, and *v* happen must be defined beforehand.

**Figure 2.**
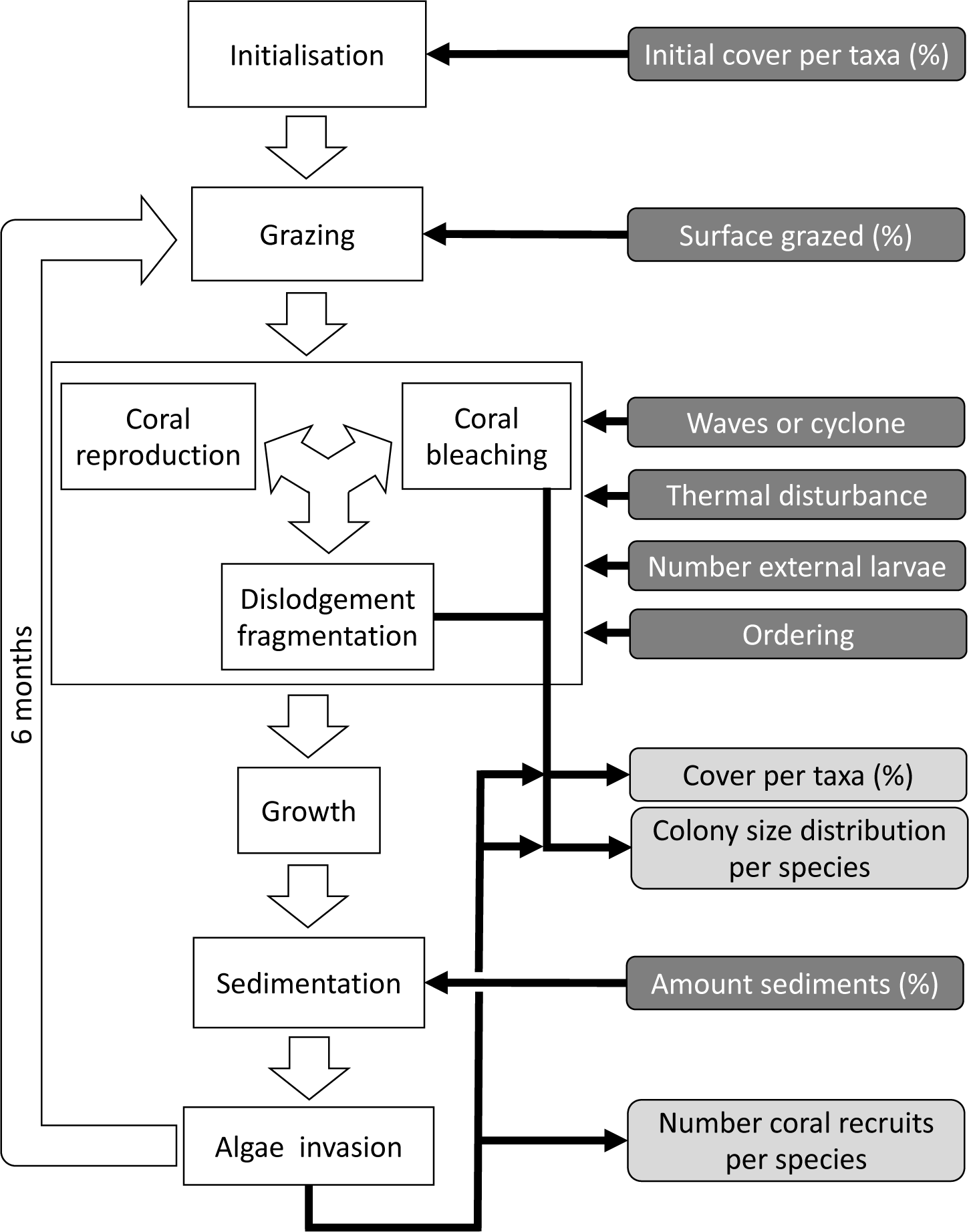
Ordering of processes in the coral agent-based model: white rectangles represent processes, dark grey rectangles with white text are input data, and light grey rectangles with black text are outputs. Large white arrows define the ordering of processes and black arrows show the direction of data transfer. The order of occurrence of coral reproduction, bleaching, and colony dislodgement and fragmentation is imposed to simulate recruitment failure due to the occurrence of a disturbance prior to reproduction. The intensity of waves and cyclones is expressed as a dimensionless dislodgment mechanical threshold; thermal stress is expressed in degree-heating weeks.

During each time step, the model exports three response variables: the cover of each benthic group (coral, algae, and sand), the planar area of each colony present per species and the number of recruited coral larvae m^-2^ species^-1^. The first two variables are collected after processes *iii, iv*, and *vii*, and the third variable after process *vii*. There are six variables imported each time step; their values respectively determine the cover to be grazed, the intensity of waves or cyclone, and of thermal stress, the number of external larvae m^-2^ entering the reef, the order that reproduction, bleaching, and wave or cyclone events happen, and the cover of sand to be achieved.

#### 2.2.4. Design and concept

##### Emergence

The dynamics of the benthic community emerge from species traits and the imposed disturbance regime (waves, cyclone, thermal stress), larval connectivity, and grazing pressure intensity.

##### Objective/Fitness

The goal of a living agent is to grow (i.e., to convert its neighbouring agents into its own state, Figure 1), which depends on its traits and those of the neighbouring agents (Appendix S2: §6). The fitness of a colony is determined by its traits and its size — resistance traits influence survival, recovery traits influence the capacity to reproduce vegetatively and sexually, and colony size influences resistance to waves and cyclones, spatial competition, as well as its fecundity.

##### Interactions

Agents on the edge of coral colonies and patches of algae interact with one another when competing for space. The outcome of a coral-coral interaction is determined by its specific pairwise outcome probabilities — the probability of coral-algae interactions are the same for all coral species, and algae-algae interactions result in a stand-off except when competing against crustose coralline algae. Branching and plating species also have the capacity to overtop other colonies and algae depending on their size (Appendix S2: §6).

##### Stochasticity

The model draws success or failure outcomes each time a patch of algae is grazed, a larva attempts to settle and survive the first six months, an agent tries to convert another leaving agent (Appendix S2: §2, 3, 6), and a colony is thermally stressed (i.e., bleaching and bleaching-induced mortality; Appendix S4: §2, 3). Each of these random events is based on a probability of success specific to the process and the species involved. During initialization, colonies are created and placed randomly in space; their sizes are drawn from right-skewed frequency distributions (Appendix S2: §1). Finally, grazing and larval settlement happen randomly in space.

##### Collectives

Collective behaviour of agents happens when a colony is (*i*) dislodged — agents sharing the same colony (i.e., coral and algae agents growing on a colony) are converted to barren ground (Appendix S2: §4), (*ii*) bleaches or dies — the coral agents of the colony are converted to a bleached or dead state, respectively (Appendix S4: §2, 3), (*iii*) reproduces — the number of larvae or gametes produces by a colony depends on certain coral traits and the size and age of the colony (Appendix S2: §3).

##### Observation

Three types of data are collected during a simulation: (*i*) percentage cover of each taxon, (*ii*) number of recruits for each coral species m^-2^, and (*iii*) planar area of each colony species^-1^ (Figure 2).

#### 2.2.5. Initialization

The initial composition of the benthic community (i.e., the cover of coral species, algae, barren ground and sand) is defined by the user and is imported from a comma-delimited text file. The space is filled by creating colonies and patches of algae randomly, then by converting the remaining agents into barren ground and sand (Appendix S2: §1).

#### 2.2.6. Input

Predefined time series (recorded in the text files) of input data are used to define the environmental context of the reef (Figure 2). At each time step, the model imports values for the corresponding period of the (*i*) surface grazed (%), (*ii*) number of external larvae settling, (*iii*) intensity of waves of cyclones (in dislodgement mechanical threshold, a dimensionless measure of the mechanical threshold imposed by waves and cyclones) and (*vi*) thermal stress intensity (in degree-heating weeks), and (*v*) sand cover (%).

#### 2.2.7. Sub-models

We defined sub-models to simulate the following processes:

##### Response to waves and cyclones

We modelled colony dislodgment using Madin and Connolly’s 2006 colony shape factor, which is compared for each colony to the intensity of the disturbance (expressed as dislodgement mechanical threshold). We implemented branching-colony fragmentation by modifying the relationship between fragment size and survival established by Highsmith et al. (1980) (Appendix S2: §4.1). We defined our own models to simulate the effect on the algae community because no relationships have been established empirically (Appendix S2: §4.2).

##### Bleaching

We first defined species-specific index of bleaching susceptibility using bleaching resistance traits and Swain and colleagues’ (2016) bleaching response index. We then used this index to establish species-specific logistic bleaching responses as a function of the intensity of the thermal stress (in degree heating-week). Finally, we defined a bleaching-induced mortality logistic-response model (Appendix S4).

##### Larval production

We defined logistic models to determine the proportion of mature polyps present in a colony, as a function of its planar area and species maximum colony diameter. We built these models using the empirically defined relationships of Álvarez-Noriega et al. (2016) (Appendix S2: §3.1.1.b). We used the models of Figueiredo et al. (2013) to determine the proportion of larvae remaining on the reef as a function of retention time (Appendix S2: §3.1.1.d). We used the models of Connolly and Baird (2010) to determine the proportion of viable and competent larvae coming from a source reef as a function of the distance travelled (Appendix S4: §3.1.2.b).

### 2.3. Model calibration

We provide here a short description of the calibration. All details are presented in Appendix S3.

#### 2.3.1. Study sites and related data

We used data collected between November 2001 and July 2011 in three sites located in Martinique in the Caribbean: Fond Boucher (14° 39’ 21.07” N, 61° 09’ 38.98” W), Pointe Borgnesse (14° 26’ 48.74” N, 60° 54’ 12.72” W), and Ilet à Rats (14° 40’ 58.04” N, 60° 54’ 1.18” W). The data were collected biannually by the *Observatoire du Milieu Marin Martiniquais* for the program *Initiative Française pour les REcifs COralliens*. These data describe the benthic, macroinvertebrate, and fish communities at the species or genus levels, as well as sand cover for each site and at each sampling time (Appendix S3: Figure S1, S2). We downloaded values of degree-heating weeks for the corresponding location from the US National Oceanic and Atmospheric Administration data server ERDDAP (Environmental Research Division’s Data Access Program; coastwatch.pfeg.noaa.gov/erddap) (Appendix S3: Figure S3). We identified cyclone tracks using the National Oceanic and Atmospheric Administration Historical Hurricane Tracks website (coast.noaa.gov/hurricanes).

#### 2.3.2. Definition of the environmental context

We modelled thermal stress by inputting at each time step the maximum degree-heating week value found for the corresponding period. We represented the intensity of hydrodynamic regimes by inputting values of the dislodgement mechanical threshold (Appendix S2; Madin and Connolly 2006). We imposed a constant value in the absence of cyclone and a lower value when Hurricane Dean affected the reefs in August 2007 (its intensity changed from Category 1 to 2 while passing over Martinique). We chose threshold values arbitrarily considering wave exposure and cyclone intensity. Because of this uncertainty, we defined three different hydrodynamic regimes that we included in the calibration procedure for each site (Appendix S3: Figure S5).

To estimate the percentage of the reef grazed at each time step, we first defined models predicting grazing intensity (i.e., percentage cover maintained in a cropped state) as a function of herbivorous fish and urchin density. We defined these models using the empirical data from Williams and Polunin (2001) and Sammarco (1980) for fish and urchins, respectively (Appendix S3: Figure S6). We then used these models and the population densities of *Acanthuridae* spp., *Scaridae* spp., and sea urchins measured in the three sites to predict their respective grazing regimes. Finally, we defined three additional similar regimes of different intensities, which we included in the calibration procedure (Appendix S3: Figure S8).

The model adjusts the amount of sand cover (i.e., by removing or adding sand patches) at each time step according to the observed cover measured in each site (Appendix S3: Figure S4). Having no information about larval connectivity at the three sites, we set the number of larvae m^-2^ at 700 during each reproductive time period (i.e., once a year). This number corresponds to our estimate of competent larvae arriving on a hypothetical reef 20 km from an upstream reef having a 50% coral cover (Appendix S2: §3.1.2). The number is realistic considering that the distance separating the three sites from other coral communities is lower, but the average coral cover in the West French Indies is on average < 40% (Wilkinson, 2008).

#### 2.3.3. General procedure

We calibrated the model for each site independently. We selected twelve parameters, for which we defined between two to five potential values (Appendix S3: Table S1). We defined an algorithm to explore the parameter space optimally. The algorithm first selects the centroid, the most extreme values, and the values situated at mid-distance between the centroid and the extremes. A simulation with each parameter value is launched and replicated five times. We measured the fit between the empirical and simulated cover time series using an objective function. The objective function measures the performance of a given run by calculating the Euclidian distance between the empirical and simulated cover time series (averaged over five replicates), averaged over all the taxa (Appendix S3: §3.2). Performance is positive, with smaller values indicating higher performance (lower difference between simulated and empirical values). The algorithm then selects the ten runs providing the best performance and generates for each of them, the five closest (using the Gower’s distance metric; Gower 1971) and untested parameter combinations. The algorithm then launches these new simulations and repeats the procedure once more.

To compare the performance of model runs to a null expectation, we generated a null distribution of performance values for each empirical dataset by randomizing cover values within each row and calculating the distance from the original datasets.

### 2.4. Hierarchically structured model validation

Models are often validated by comparing outputs of a single level of organization (i.e., individual, population, community) to equivalent empirical datasets (individual species covers in our case), but this approach only examines lower dimensionality for more complex models. Following the recommendation of Kubicek et al. (2015), and aligned with the approach of pattern-oriented modelling (Grimm et al., 2005), we assessed whether the different processes implemented in our model produce ecologically realistic patterns by comparing them to expectations formulated *a priori*. We assessed the following processes of our model: (*i*) we expected colony lateral growth to equal the species growth rate in absence of spatial interaction and to decrease as space becomes saturated by colonies; (*ii*) recruitment rate should increase as a population grows from low initial cover, and then decreases as space saturates; for competition under different (*iii*) disturbance-regime intensities — we expected the competitive species to dominate the community under low-disturbance regimes, and ruderal or stress-tolerant species otherwise; (*iv*) larval connectivity — we expected species with higher colony fecundity or brooding mode of larval development to dominate the community under low connectivity, and the competitive ones otherwise; (*v*) grazing — under low grazing pressure, the benthic community should be dominated by algae, and by corals otherwise. Because we expected the community dynamics to depend on species-specific trait differences, we did procedures *iii, iv*, and *v* with two different communities, each composed of a competitive, a ruderal, and a stress-tolerant species, originating from the Eastern Pacific and Western Atlantic, respectively. Note that our goal was not to use suites of species that accurately reflect the taxonomic composition of particular reefs or species pools, but rather to select species based on their functional trait attributes. Thus, although we refer to species by name, the names themselves matter less than their functionality. It is well known that reefs with different biogeographic or evolutionary histories host species that are functionally very similar (McWilliam et al., 2018b). All details are in Appendix S5.

### 2.5. Global sensitivity analysis

We constructed a global sensitivity analysis for 10 of the calibrated parameters and six additional parameters with high uncertainty (Appendix S6: Table S1). For each parameter, we defined a range around the value(s) calibrated (for the 10 parameters considered in the calibration) or the value used in the simulations (for the six additional parameters). These parameters are all continuous but vary in their type (i.e., probabilities, ratio, heights, sub-model coefficients). We defined their respective ranges considering parameter uncertainty, realistic boundaries, and what values might improve the model performance based on the model calibration and hierarchically structured validation. We did the procedure for each site independently because certain parameters were calibrated on different values between sites and because the coral communities differ.

We simulated our model for 10 years with a bleaching event of an intensity of 12 degree-heating weeks occurring after four years. We kept the following processes constant: grazing (50%), wave hydrodynamic regime (dislodgement mechanical threshold = 120), and larval input from regional pool (700 larvae m^-2^). We defined the same initial benthic composition as the one observed in the Caribbean sites.

We defined five response variables that represent the ecological state of the community at the end of the simulation: (*i*) total coral cover, (*ii*) difference of total coral cover at year 10 and just after the bleaching event, (*iii*) Pielou’s evenness, (*iv*) coral species richness (only the species having ≥ 1% cover), and (*v*) number of recruits m^-2^.

We estimated the relative importance of the parameters selected on each response variable following the efficient protocol of Prowse et al. (2016). For each site, we sampled 1000 combinations of parameter values from a continuous parameter space using Latin hypercube sampling and continuous distributions. We launched each combination once (no replicates). We then fitted boosted regression trees on the input parameter values for each response variable — the procedure provides the respective influence of each predictor (i.e., model parameter) on the variation of the response variable. We ensured the sampling was sufficient by comparing the influence of the parameters obtained with *n* = 1000 samples with values obtained with subsamples (*n* = 100, 250, 500 and 750); sampling is estimated sufficient when the influence of the parameters converge to similar values as sample size increases. All the details of the procedure are in Appendix S6.

## 3. Results

### 3.1. Model calibration

Model performance varies between 28 and 10 (lower values = better performance), and were all lower than the lower 95% confidence bound of the random distribution (Appendix S3: Figure S9). This shows that despite the model’s complexity and parameter uncertainty, the model outputs population dynamics closer to the empirical data compared to random. The best performance values converged toward 10 among the three sites (i.e., minimum ± standard error: 10.93 ± 3.677, 10.89 ± 2.872, 10.39 ± 3.119 for Fond Boucher, Pointe Borgnesse and Ilet à Rats, respectively).

With the combination of parameter estimates yielding the best fit, the model produces time series of total coral cover similar to the empirical ones for each site (Figure 3; Appendix S3: Figure S15, S16). The difference between the simulated and real total coral cover does not exceed 15, 20, and 11% for Fond Boucher, Pointe Borgnesse and Ilet à Rats, respectively. Results at the species level are more variable, but the cover difference of individual coral populations never exceeds 8%. For some species, the simulated cover closely predicts the empirical data — for instance, *O. faveolata* and *O. annularis* at Ilet à Rats (Figure 3) and *A. agaricites* and *S. siderea* at Fond Boucher (Appendix S3: Figure S15). The model failed to predict the population dynamics of some other species accurately; for instance, in the simulated reefs, *M. mirabilis, M. decactis* and *P. furcata* became the dominant species, while the cover of *P. atreoides* and *M. meandrites* approached zero at Fond Boucher; *M. mirabilis* outcompeted *O. annularis, O. faveolata, O. franksi* and *P. astreoides* at Pointe Borgnesse (Appendix S3: Figure S15, S16), while the *P. astreoides*’s population decreased at Ilet à Rats (Figure 3).

**Figure 3.**
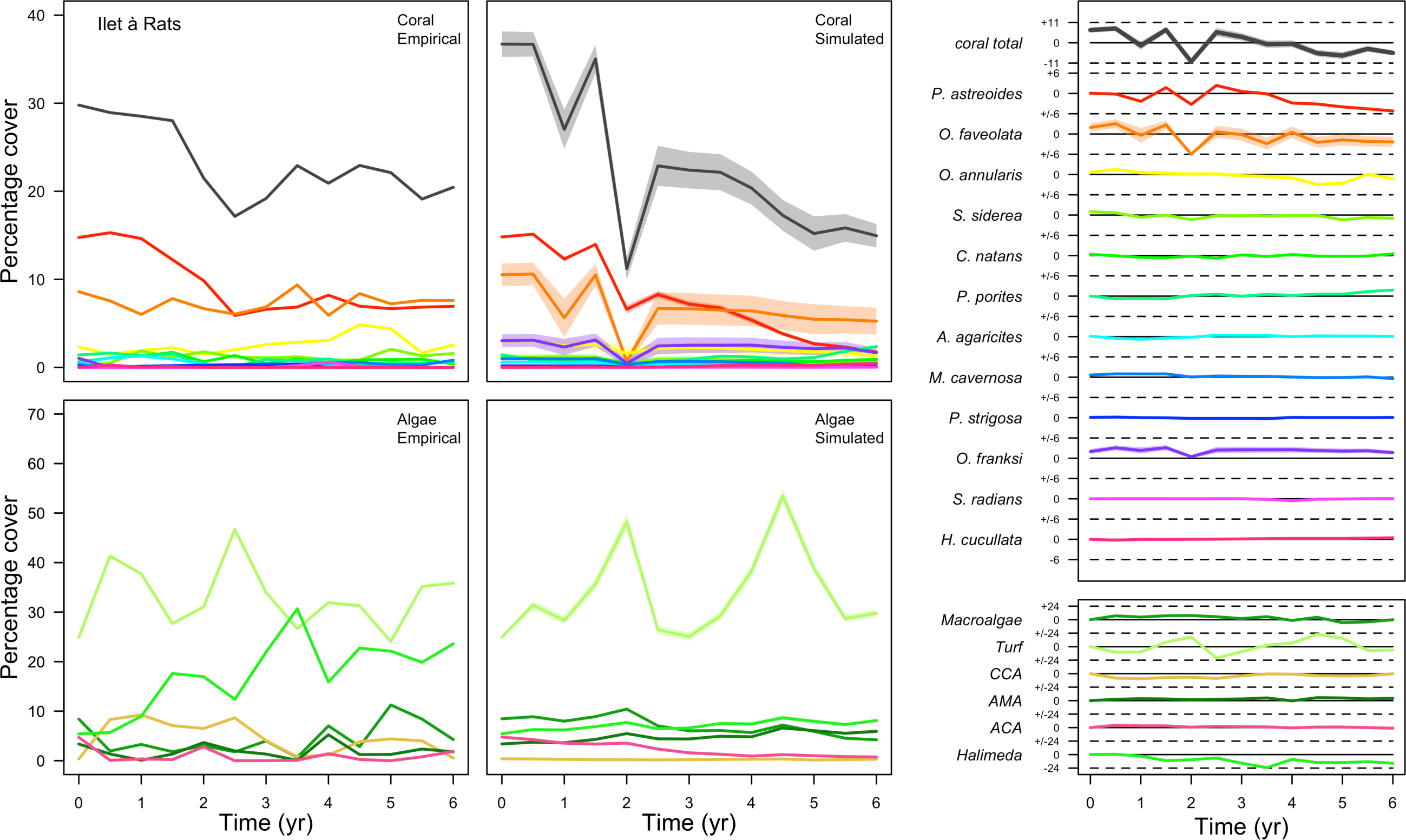
Comparison of empirical and simulated taxa cover for the combination of parameter values providing the best fit for site Ilet à Rats. Solid lines in the simulated time series are the mean percentages cover (averaged over five replicates) and the shaded areas show the standard error. The right panels display the cover difference between simulated and empirical time series.

The simulated cover of algae also closely mimics the empirical data for most algal groups (Figure 3, Appendix S3: Figure S15, S16). The difference of percentage cover is the highest for turf and reaches a maximum of 29, 22 and 24% for Fond Boucher, Pointe Borgnesse and Ilet à Rats, respectively. These percentages are high compared to other groups or taxa, but this can be explained partially by the high variance in algal turf cover observed at the reefs. Turf cover generally fluctuates by > 20%, a pattern that our model was able to reproduce at all three sites (Figure 3; Appendix S3: Figure S15, S16).

Notably, crustose coralline algae are systematically less abundant in the simulated reefs compared to the observed data, a phenomenon we attempted to correct in the calibration procedure (Appendix S3: §3.3). Finally, the model could not reproduce the high cover of *Halimeda* spp. observed at Ilet à Rats (Figure 3) compared to the other sites. See Appendix S3 for more detailed results and discussion regarding the between-site comparison.

### 3.2. Hierarchically structured validation

The hierarchically structured validation shows that the model produces ecologically realistic population dynamics under different environmental conditions. Here we provide a summary of the results, but a more-complete description and explanation are available in Appendix S5.

#### Growth

As expected, coral colonies grew at their species-specific growth rate at low population density. However, as space filled up, colonies began constraining each other spatially and their growth rates decreased until eventual stasis (Appendix S5: Figure S4).

#### Recruitment

For a single coral population, the different patterns of recruitment observed among three functionally distinct species (Appendix S5: Figure S6) results from the interaction of several factors: (*i*) individual colony fecundity determined by its planar area, species-specific polyp fecundity, corallite area (polyp size), growth form, sexual system, and mode of larval development, (*ii*) the distribution of colony size in the population, and (*iii*) the amount of surface available for larval settlement. Weedy (*Agaricia tenuifolia*) and stress-tolerant species (*Echinophyllia orpheensis*) produced bell-shape recruitment patterns (Appendix S5: Figure S8). Recruitment rate was initially low because populations were composed of small, low-fecundity colonies, but the rate increased as colonies grew and became more fertile (Appendix S5: Figure S8). Recruitment subsequently decreased as space became saturated. In contrast, recruitment rate for competitive species (*Acropora gemmifera*) was initially high and only decreased as cover occupancy increased. This pattern is essentially due to a higher vegetative growth rate associated with a population initially composed of fewer but larger, more fecund colonies (Appendix S5: Figure S8).

#### Disturbance intensity

In both the Western Atlantic and Eastern Pacific communities, the competitive species dominated the coral community under low wave exposure (Appendix S5: Figure S10, S11, S14). The success of the competitive species was due mainly to two interacting processes — with a higher vegetative growth rate, competitive species (*i*) overcame free space before other species, and (*ii*) enhanced recruitment by achieving large colony size rapidly. Higher wave exposure reduced the cover of competitive species because colonies were dislodged at a certain colony size, which reduced recruitment rate and provided other species with more available space to grow and recruit.

In the Western Atlantic community, increased availability of space favoured the weedy species (*Madracis pharencis*) over the stress-tolerant species (*Orbicella annularis*), principally because of the former’s brooding mode of larval development (twice a year), faster growth rate, and high wave-resistance of its growth form (digitate). In contrast, species coexisted in the Eastern Pacific community under the highest-intensity disturbances (Appendix S5: Figure S14). Both the competitive (*Pocillopora elegans*) and stress-tolerant species (*Porites lutea*) recruited more than weedy species (*P. damicornis*) due to their spawning mode of reproduction; spawning species received three times more larvae from the regional pool (Appendix S2: §3.1.2). However, the weedy species recruited twice as frequently and is slightly more aggressive than the other two species. The stress-tolerant species has a massive growth form, which conferred higher resistance to waves compared to the other two branching species. Nonetheless, this advantage barely compensated for its slower growth rate and lower colony fecundity.

Population(s) recovered to pre-disturbance cover only after one year, regardless of the intensity of the event. This recovery is faster than most dynamics observed in real reef systems and arises because we imposed a constant and high number (7000 larvae m^-2^) of larvae coming from the regional pool. In reality, larval supplies are reduced because a strong bleaching disturbance would also affect the surrounding reefs (Hughes et al., 2019). Another reason for this outcome was that recruitment preceded growth in the model (Figure 2), which inflated the former process because more space was available for settlement.

#### Larval connectivity

Low larval connectivity influenced the two coral communities differently (Appendix S5: §5). In the Western Atlantic, the weedy species thrived under zero to moderate larval input (0, 66, 700 larvae m^-2^) while the other two species went locally extinct (Appendix S5: Figure S17). The weedy species produced ready-to-settle larvae twice a year, while the other two species reproduced annually and only a portion of their larvae were able to settle because of their time to motility (Appendix S2: §3.1.2.1.d). In contrast, the stress-tolerant species dominated in the Eastern Pacific community (Appendix S5: Figure S18), due to its higher wave-resistance compared to the other two branching species.

Under the highest larval connectivity (7000 and 35,000 larvae m^-2^), the competitive species dominated in both communities, principally because of their higher growth rates, spawning mode of reproduction, and their capacity to overtop smaller colonies.

#### Grazing

Population dynamics were similar between the two communities (Appendix S5: Figure S20, S25). As expected, the percentage of reef grazed corresponded approximatively to the total coral cover at the steady state; the remaining ungrazed part of the reef was occupied by algae. We observed no hysteresis because we did not implement feedback processes. Turf dominated the algae community in all simulated grazing regimes, despite having the highest palatability among algae (Appendix S2: Table S2). The success of this species was due to its much higher growth rate compared to other algae (Appendix S2: Table S17).

Coral recruitment rates at the steady state were the highest under medium grazing pressure (50%) because under lower and higher grazing intensities, space was saturated by turf and coral colonies, respectively. Under the lowest grazing pressure, most of the colonies were ≤ 100 cm^2^ in surface area (Appendix S5: Figure S22) and coral populations were rescued by external larval input. At intermediate pressures (30 and 50%), the competitive species dominated the coral community mainly because of their higher external larval input compared to the brooding (weedy) species, and their higher growth rate than the stress-tolerant species. Having a high growth rate was particularly important under low grazing pressure because this trait compensated better for the cover lost in the competition with turf, which won all its interactions with corals in the model (Appendix S2: Table S17).

Under higher grazing pressures (70 and 90%), there were more coral-coral and fewer coral-algae interactions, which changed coral species dominance. The Western Atlantic community was dominated by the weedy species, followed by the tress-tolerant species and the competitive species was competitively excluded, mainly because of its lower aggressiveness and highest vulnerability against waves (Appendix S5: Figure S10). In the Eastern Pacific community, the stress-tolerant species dominated the coral community and slowly outcompeted the other two species principally because of its much higher wave resistance (Appendix S5: Figure S27).

### 3.3. Global sensitivity analysis

The parameters that had the most important effects on the response variables were *growth rate reduction interaction* (the reduction of lateral growth rate of an organism overgrowing another one) and *otherProportions* (coefficient controlling the number of larvae produced locally), followed by probabilities for larvae to settle on different substrates, and the probabilities of algal grazing. The remaining ten parameters did not have an important influence on any of the five response variables (Appendix S6: Figure S1).

Globally, all the influential parameters affected the response variables according to expectations. For instance, increasing growth rate reduction when organisms interact (mostly turf over corals) reduced the competitive advantage of the dominant taxa, which increased coral *richness, total coral cover* (due to reduced competitiveness of algae), and consequently enhanced the *difference in coral cover* and the *number of coral recruits* (Appendix S6: Figure S1). Increasing *otherProportions* increased the number of larvae produced by each coral population, which positively affected *coral cover, cover difference* and the *number of coral recruits*. The parameter was negatively correlated with *richness* and *evenness* because higher values disproportionately benefitted species capable of higher recruitment (e.g., brooder; Appendix S6: Figure S3).

In general, the parameters influenced the response variables in consistent ways among sites (Appendix S6: Figure S1). Differences were mainly due to different ranges of values tested for particular parameters. For instance, *probability of grazing allopathic macroalgae* had a stronger effect at Pointe Borgnesse compared to the other two sites because its range included smaller values, implying lower palatability, higher abundance (Appendix S6: Table S1, Figure S4), and a higher effect.

Ten of the parameters had a negligible effect on the response variables, because the processes they contributed to did not occur in these simulations. For instance, *probability of algae to cover crustose coralline algae* did not have an effect because crustose coralline algae was not present in high enough abundance (Appendix S6: Figure S4), and *height of big algae* and *height of turf* did not have an effect because most of the branching colonies did not reach sizes large enough to overtop these algae (Appendix S6: Figure S5).

## 4. Discussion

Our primary goal was to develop a model that captured the spatio-temporal dynamics of community composition in coral reefs as component coral and algal species responded to inter-species competitive interactions and external disturbances. Our trait-based and demographic approaches provided a combination that yielded better predictions and a better understanding of coral ecosystem dynamics relative to single-component models (Edmunds et al., 2014; Salguero-Gómez et al., 2018; Violle et al., 2007). The spatial structure we imposed — a grid of 1 cm^2^ agents that collectively comprise a sizeable reef (tens of m^2^) as inspired by previous models (e.g., Langmead and Sheppard 2004, Sleeman et al. 2005, Tam and Ang 2009, Sandin and McNamara 2012) — yielded emergence, scaling, self-organization, and unpredictability, each of which is a property of complex systems (Parrott, 2002) including coral reefs (Dizon and Yap, 2006; Hatcher, 1997). Operating at such a small spatial grain, processes can be modelled at the appropriate scale (e.g., dislodgement removes entire colonies while spatial competition affects colony edges) (Figure 1) to generate distributions of colony size, and in turn, colony fitness and performance. Overall, the population dynamics resulted from the collective performance of each colony, which implies that at the scale of the community, a given species’ fitness depended on its capacity to persist under a certain environmental context and compete with functionally dissimilar species. As in the real world, macro-scale community dynamics emerged from finer-scale processes and interactions, a phenomenon clearly demonstrated by the hierarchically structured validation (Appendix S5). The model structure can also accommodate realistic spatial colony arrangement — a feature that is absent in previous models (but see Wakeford et al. 2008), despite the importance of spatial patterns for herbivory (Eynaud et al., 2016) and coral population dynamics (Brito-Millán et al., 2019).

Our model is unique in being designed to simulate the effects of coral species richness and functional diversity on ecosystem dynamics. Most coral models have been developed to describe the effect of external drivers (mainly disturbances) on the state of the coral community (usually total cover) (e.g., Melbourne-Thomas et al. 2011a, Madin et al. 2012b, Bozec and Mumby 2015, Kubicek and Reuter 2016). In contrast, few models exist that assess the influences of aspects of diversity on community or ecosystem dynamics (e.g., Tam and Ang 2012, Ortiz et al. 2014, Fabina et al. 2015), but these have represented diversity with limited detail, and consequently have little capacity to evaluate the effects of identity and diversity on ecosystem functioning (Brandl et al., 2019), or the effects of functional redundancy and response diversity on ecosystem resilience (Mcleod et al., 2019). In contrast, our model represents diversity in detail — we considered eleven functional traits and included their influence over eight ecological processes applied to 798 functionally realistic species. In addition, we ensured that species richness can be varied without affecting computation time. Our model therefore enables exploration of many realistic assemblage scenarios within an easily modified experimental setting.

Our calibrated model was able to reproduce similar total coral cover dynamics in the three sites. At the population level, results were more varied, with several populations well predicted, others less so. Overall these results are remarkable considering model complexity, the high number of parameters, and the limited data describing the environmental context and diversity at the three sites.

A useful model should ideally be calibrated and validated with empirical data at each level of organization (e.g., colony, population, community) (Kubicek et al., 2015). However, empirical data are usually lacking for some or even all of these levels. Coral models have therefore been validated against one or a few community-aggregated variables, and rarely at the species level. Sampling additional data specifically for the model and at the sites used for calibration and benefiting from the opinion of local experts can improve the capacity of the model considerably to reproduce realistic dynamics. For instance (Mumby, 2006) developed a spatially explicit mechanistic model to reproduce the total coral and macroalgae cover observed in Jamaican reefs. In two subsequent developments of the model, Ortiz et al. (2014) reproduced accurate recovery rates and final community composition of six coral taxa at fourteen reefs in the Great Barrier Reef, and Bozec et al. (2015) reproduced the cover of seven coral species and the rugosity in reefs in Cozumel (Mexico). With a similar model, Kubicek et al. (2012) generated time series of major coral taxa cover at Chumbe Island (Tanzania) similar to real data. Further, Kayal et al. (2018) accurately reproduced colony density distributions of three coral species in four different sites in Moorea, French Polynesia, using integral-projection models.

In contrast with conventional approaches to model development, we developed our model independently of the empirical data upon which calibration was based. Instead, our model included the ecological details required for achieving our primary objective. Below we discuss the primary sources of uncertainty in our model calibration and suggest realistic ways for improvement.

### Grazing

We estimated average grazing pressure over six months (% of reef grazed) based on a biannual assessment of sea urchin and herbivorous fish (*Scaridae* spp. and *Acanthuridae* spp.) populations. *Acanthuridae* spp. are mobile herbivores (Thibaut et al., 2012), so frequent assessments are necessary to obtain accurate estimates of mean population size. More data collection could improve the accuracy of the modelled processes, has others have done for several fish species (e.g., Bozec et al., 2016; Mumby, 2006).

### Hydrodynamic regime

We defined time series of dislodgement mechanical threshold as a function of site exposure and cyclone intensity. Measuring the real dislodgement mechanical threshold over time in each site would improve the precision of the simulations. This would require measuring horizontal water velocity and tensile strength of the substratum (Madin et al., 2012a; Madin and Connolly, 2006).

### Recruitment

There is high uncertainty in our implementation of recruitment because we did not have estimates of recruitment rates and of the proportion of recruits originating from the local reef *versus* the regional pool. We therefore fixed the number of external larvae coming into the reef and controlled recruitment rate with one parameter (*otherProportion*s). The sensitivity analysis revealed that the parameter has a strong influence on the model’s predictions. Reducing this uncertainty requires better estimates of recruitment rates, which can be achieved with tile experiments (e.g., Ritson-Williams et al. 2016) or visual assessment of new recruits along transects (e.g., Gilmour et al., 2013; Holbrook et al., 2018). The proportion of locally *versus* regionally recruited larvae can be estimated with population genetics (e.g., Almany et al., 2017; Johnson et al., 2018) or by modelling larvae plumes (e.g., Golbuu et al., 2012; Wolanski and Kingsford, 2014).

### Trait data

The hierarchically structured validation showed that between-species trait differences influenced community dynamics. Considerable gaps in the coral-trait database (Madin et al., 2016a) limited our capacity to estimate traits accurately for many coral species. Collecting reliable trait data is critical to predict coral-community dynamics and ecosystem functioning (Madin et al., 2016b). Further precision in the prediction would be gained by measuring traits locally, because traits can vary substantially among populations in different locations (e.g., Diaz-Pulido et al., 2009), and factors such as nutrient concentration affect both algae and corals growth (Wear and Thurber, 2015; Zaneveld et al., 2016).

Our model is flexible and can be tailored to represent coral communities around the world, and to explore many different questions pertaining to the links between diversity and ecosystem dynamics. This version of the model focusses primarily on coral diversity and the effect of two disturbance types, but other disturbance types, and additional processes and aspects of reef diversity, could be easily implemented, provided sufficient data are available. Examples include functions related to herbivory, algal diversity, disturbance types, and feedback processes, which we elaborate below.

Herbivores differ in their foraging behaviour (reviewed in Appendix S2: §2.1), which affects benthic diversity, coral reef recovery, and functioning (Burkepile and Hay, 2010; Cheal et al., 2013, 2010; Nash et al., 2016; Pratchett et al., 2014). A few models have described aspects of herbivore diversity; for instance, Sandin and McNamara (2012) modelled the effect of spatially differentiated foraging behaviour between fish and urchins on the dynamics of a coral community, and Bozec et al. (2016) modelled the population dynamics of several parrot fish species and their respective species and size-specific contribution to grazing. However, herbivore diversity has generally been neglected in coral-reef models. Accommodating herbivore diversity and its effect on the benthic community in our model is feasible, provided associations between population densities and processes (e.g., grazing, bioerosion) are empirically established for different taxonomic or functional groups (e.g., Appendix S3: §2.3).

Algal diversity is potentially as important as coral and herbivore diversity for reef functioning and recovery (e.g., Roff et al. 2015). Yet, most coral-reef models describe the algal community with no more than three functional groups (macroalgae, crustose coralline algae, turf). Our model is the first to implement six functional groups, which accommodated additional ecological details such as grazing preferences and coral-algae interactions (Appendix S2: §2.1 and §6.3, respectively). To date, trait-based research on tropical reef algae is modest compared to fishes and corals (Brandl et al., 2019) and an algal-traits database has not yet been created.

We implemented the effects of hydrodynamic variation, thermal disturbances, and changes in grazing pressure, but reefs are also affected by other disturbances, and some of these have been implemented in previous models — including ocean acidification (e.g., Anthony et al., 2011; Madin et al., 2012b), predation by *Acanthaster planci* (e.g., Hogeweg and Hesper, 1990; Van der Laanm and Bradbury, 1990), disease (e.g., Brandt and McManus, 2009), destructive fishing (e.g., Kubicek et al., 2012), and pollution (e.g., Wolanski et al. 2004, Melbourne-Thomas et al. 2011, Kennedy et al. 2013). We are currently not able to model the species-specific effects of these disturbances on coral assemblages because it is not clear what traits are relevant, nor how these relate to ecological processes and responses. Such information is necessary to parameterize mechanistic models such as ours, as exemplified by our trait-based model of the response of corals to bleaching (Appendix S4). Nevertheless, our model would benefit from further validation, and is missing important variables (e.g., symbiont diversity) for which data are lacking (Carturan et al. 2018).

Feedback processes affect population dynamics by generating thresholds, hysteresis, and by shaping basins of attraction (Scheffer et al., 2001; Scheffer and Carpenter, 2003). Coral reefs are notorious for feedback processes (Hughes et al., 2010; Mumby and Steneck, 2008), some of which have been implemented in models (e.g., Mumby et al. 2007, Muthukrishnan et al. 2016, Kubicek and Reuter 2016). Leemput et al. (2016) reviewed over 20 different feedbacks processes observed in reefs and demonstrated with a simple model that the combination of several feedback processes, although weak individually, can have important effects on system dynamics. However, the empirical quantification of these processes remains to be established (van de Leemput et al., 2016).

## 5. Conclusion

We have constructed a dynamic and customizable model that allows coral species richness and functional diversity to be manipulated independently. The model combines trait-based, demographic and complex system science approaches to implement many ecological processes that drive coral reef dynamics. Its structure is flexible, and more processes, traits and taxa can be incorporated, provided the data are available. To that end, we highlighted several knowledge gaps that impede the modeling of important details or components of coral reef ecosystems. Our model can be used as a platform for virtual experiments aimed at testing hypotheses about the effects of species identity and diversity on ecosystem functioning, and about the effects of functional redundancy and response diversity on resilience.

## Acknowledgements

Constructive feedback from M. Julian Caley, Kristen T. Brown and Sun W. Kim and technical support from Wade Klaver and Fabian Cid Yañez is gratefully acknowledged.

## Competing interests

The authors declare that no competing interests exist.

## Supporting information

http://dx.doi.org/10.17605/OSF.IO/CTQ43

## References

Almany GR, Planes S, Thorrold SR, Berumen ML, Bode M, Saenz-Agudelo P, Bonin MC, Frisch AJ, Harrison HB, Messmer V, Nanninga GB, Priest MA, Srinivasan M, Sinclair-Taylor T, Williamson DH, Jones GP. 2017. Larval fish dispersal in a coral-reef seascape. Nat Ecol Evol 1:1–7. doi:10.1038/s41559-017-0148

Alvarez-Filip L, Carricart-Ganivet JP, Horta-Puga G, Iglesias-Prieto R. 2013. Shifts in coral-assemblage composition do not ensure persistence of reef functionality. Sci Rep 3:3486. doi:10.1038/srep03486

Alvarez-Filip L, Dulvy NK, Gill JA, Côté IM, Watkinson AR. 2009. Flattening of Caribbean coral reefs: region-wide declines in architectural complexity. Proc R Soc B Biol Sci 276:3019–3025. doi:10.1098/rspb.2009.0339

Álvarez-Noriega M, Baird AH, Dornelas M, Madin JS, Cumbo VR, Connolly SR. 2016. Fecundity and the demographic strategies of coral morphologies. Ecology 97:3485–3493. doi:10.1002/ecy.1588

Anthony KRN, Maynard JA, Diaz-Pulido G, Mumby PJ, Marshall PA, Cao L, Hoegh-Guldberg O. 2011. Ocean acidification and warming will lower coral reef resilience. Glob Chang Biol 17:1798–1808. doi:10.1111/j.1365-2486.2010.02364.x

Bartón K. 2017. MuMIn: multi-model inference. R Packag version 1.40.0.

Bates D, Mächler M, Bolker B, Walker S. 2015. Fitting linear mixed-effects models using lme4. J statis 67:1–48. doi:10.18637/jss.v067.i01

Bellwood DR, Streit RP, Brandl SJ, Tebbett SB. 2019. The meaning of the term ‘function’ in ecology: A coral reef perspective. Funct Ecol 33:948–961. doi:10.1111/1365-2435.13265

Bozec Y-M, Alvarez-Filip L, Mumby PJ. 2015. The dynamics of architectural complexity on coral reefs under climate change. Glob Chang Biol 21:223–235. doi:10.1111/gcb.12698

Bozec Y-M, Mumby PJ. 2015. Synergistic impacts of global warming on the resilience of coral reefs. Philos Trans R Soc B Biol Sci 370:20130267–20130267. doi:10.1098/rstb.2013.0267

Bozec Y-M, O’Farrell S, Bruggemann JH, Luckhurst BE, Mumby PJ. 2016. Tradeoffs between fisheries harvest and the resilience of coral reefs. Proc Natl Acad Sci 113:201601529. doi:10.1073/pnas.1601529113

Brandl SJ, Rasher DB, Côté IM, Casey JM, Darling ES, Lefcheck JS, Duffy JE. 2019. Coral reef ecosystem functioning: eight core processes and the role of biodiversity. Front Ecol Environ 17:445–454. doi:10.1002/fee.2088

Brandt ME, McManus JW. 2009. Dynamics and impact of the coral disease white plague: insights from a simulation model. Dis Aquat Organ 87:117–33. doi:10.3354/dao02137

Brito-Millán M, Werner BT, Sandin SA, Mcnamara DE, Brito-milla M. 2019. Influence of aggregation on benthic coral reef spatio-temporal dynamics. R Soc Open Sci 6. doi:10.1098/rsos.181703

Bulleri F, Bruno JF, Silliman BR, Stachowicz JJ. 2016. Facilitation and the niche: Implications for coexistence, range shifts and ecosystem functioning. Funct Ecol 30:70–78. doi:10.1111/1365-2435.12528

Burkepile DE, Hay ME. 2010. Impact of herbivore identity on algal succession and coral growth on a Caribbean reef. PLoS One 5:e8963. doi:10.1371/journal.pone.0008963

Carnell R. 2018. lhs: Latin hypercube samples. R Packag version 0.16.

Carturan BS, Parrott L, Pither J. 2018. A modified trait-based framework for assessing the resilience of ecosystem services provided by coral reef communities. Ecosphere 9:24. doi:10.1002/ecs2.2214

Cheal AJ, Emslie MJ, Aaron MM, Miller I, Sweatman H, MacNeil MA, Miller I, Sweatman H. 2013. Spatial variation in the functional characteristics of herbivorous fish communities and the resilience of coral reefs. Ecol Appl 23:174–88. doi:10.1890/11-2253.1

Cheal AJ, MacNeil MA, Cripps E, Emslie MJ, Jonker M, Schaffelke B, Sweatman H. 2010. Coral–macroalgal phase shifts or reef resilience: links with diversity and functional roles of herbivorous fishes on the Great Barrier Reef. Coral Reefs 29:1005–1015. doi:10.1007/s00338-010-0661-y

Clements CS, Hay ME. 2019. Biodiversity enhances coral growth, tissue survivorship and suppression of macroalgae. Nat Ecol Evol 3:178–182. doi:10.1038/s41559-018-0752-7

Connolly SR, Baird AH. 2010. Estimating dispersal potential for marine larvae: dynamic models applied to scleractinian corals. Ecology 91:3572–3583. doi:10.1890/10-0143.1

Cribari-Neto F, Zeileis A. 2010. Beta Regression in R. J Stat Softw 34:1–24. doi:10.18637/jss.v034.i02

Darling ES, Graham NAJ, Januchowski-Hartley FA, Nash KL, Pratchett MS, Wilson SK. 2017. Relationships between structural complexity, coral traits, and reef fish assemblages. Coral Reefs 36:561–575. doi:10.1007/s00338-017-1539-z

Diaz-Pulido G, McCook LJ, Dove S, Berkelmans R, Roff G, Kline DI, Weeks S, Evans RD, Williamson DH, Hoegh-Guldberg O. 2009. Doom and boom on a resilient reef: climate change, algal overgrowth and coral recovery. PLoS One 4. doi:10.1371/journal.pone.0005239

Dizon RT, Yap HT. 2006. Understanding coral reefs as complex systems: degradation and prospects for recovery. Sci Mar 70:219–226. doi:10.3989/scimar.2006.70n2219

Edmunds PJ, Burgess SC, Putnam HM, Baskett ML, Bramanti L, Fabina NS, Han X, Lesser MP, Madin JS, Wall CB, Yost DM, Gates RD. 2014. Evaluating the causal basis of ecological success within the scleractinia: an integral projection model approach. Mar Biol 161:2719–2734. doi:10.1007/s00227-014-2547-y

Eynaud Y, Mcnamara DE, Sandin SA. 2016. Herbivore space use influences coral reef recovery. R Soc Open Sci 3:160262. doi:10.1098/rsos.160262

Fabina NS, Baskett ML, Gross K. 2015. The differential effects of increasing frequency and magnitude of extreme events on coral populations. Ecol Appl 25:1534–1545. doi:10.1890/14-0273.1

Figueiredo J, Baird AH, Connolly SR. 2013. Synthesizing larval competence dynamics and reef-scale retention reveals a high potential for self-recruitment in corals. Ecology 94:650–659. doi:10.1890/12-0767.1

García AP, Rodríguez-Patón A. 2016. Analyzing Repast Symphony models in R with RRepast package. bioRxiv 047985. doi:10.1101/047985

Gillis LG, Bouma TJ, Jones CG, Van Katwijk MM, Nagelkerken I, Jeuken CJL, Herman PMJ, Ziegler AD. 2014. Potential for landscape-scale positive interactions among tropical marine ecosystems. Mar Ecol Prog Ser 503:289–303. doi:10.3354/meps10716

Golbuu Y, Wolanski E, Idechong JW, Victor S, Isechal AL, Oldiais NW, Idip D, Richmond RH, van Woesik R. 2012. Predicting Coral Recruitment in Palau’s Complex Reef Archipelago. PLoS One 7:1–10. doi:10.1371/journal.pone.0050998

Gower JC. 1971. A general coefficient of similarity and some of its properties. Biometrics 27:857–871. doi:10.2307/2528823

Graham NAJ, Cinner JE, Norström A V., Nyström M. 2014. Coral reefs as novel ecosystems: embracing new futures. Curr Opin Environ Sustain 7:9–14. doi:10.1016/j.cosust.2013.11.023

Graham NAJ, Nash KL. 2013. The importance of structural complexity in coral reef ecosystems. Coral Reefs 32:315–326. doi:10.1007/s00338-012-0984-y

Griffin JN, O’Gorman EJ, Emmerson MC, Jenkins SR, Klein AM, Loreau M, Symstad A. 2009. Biodiversity and the stability of ecosystem functioning In: Naeem S, Bunker DE, Hector A, Loreau M, Perrings C, editors. Biodiversity, Ecosystem Functioning, and Human Wellbeing: An Ecological and Economic Perspective. Oxford Scholarship Online. pp. 78–93. doi:10.1093/acprof:oso/9780199547951.003.0006

Grimm V, Berger U, Bastiansen F, Eliassen S, Ginot V, Giske J, Goss-Custard J, Grand T, Heinz SK, Huse G, Huth A, Jepsen JU, Jørgensen C, Mooij WM, Müller B, Pe’er G, Piou C, Railsback SF, Robbins AM, Robbins MM, Rossmanith E, Rüger N, Strand E, Souissi S, Stillman RA, Vabø R, Visser U, DeAngelis DL. 2006. A standard protocol for describing individual-based and agent-based models. Ecol Modell 198:115–126. doi:10.1016/j.ecolmodel.2006.04.023

Grimm V, Berger U, DeAngelis DL, Polhill JG, Giske J, Railsback SF. 2010. The ODD protocol: A review and first update. Ecol Modell 221:2760–2768. doi:10.1016/j.ecolmodel.2010.08.019

Grimm V, Revilla E, Berger U, Jeltsch F, Mooij WM, Railsback SF, Thulke H-H, Weiner J, Wiegand T, Deangelis DL, Grimm V, Revilla E, Berger U, Jeltsch F, Mooij WM, Railsback SF, Thulke H-H, Weiner J, Wiegand T, Deangelis DL. 2005. Pattern-oriented modeling of agent-based complex systems: lessons from ecology. Science (80-) 310:987–91. doi:10.1126/science.1116681

Harborne AR, Mumby PJ, Micheli F, Perry CT, Dahlgren CP, Holmes KE, Brumbaugh DR. 2006. The functional value of Caribbean coral reef, seagrass and mangrove habitats to ecosystem processes. Adv Mar Biol 50:57–189. doi:10.1016/S0065-2881(05)50002-6

Harris DL, Rovere A, Casella E, Power H, Canavesio R, Collin A, Pomeroy A, Webster JM, Parravicini V. 2018. Coral reef structural complexity provides important coastal protection from waves under rising sea levels. Sci Adv 4:1–8. doi:10.1126/sciadv.aao4350

Hatcher BG. 1997. Coral reef ecosystems: how much greater is the whole than the sum of the parts? Coral Reefs 16:S77–S91. doi:10.1007/s003380050244

Highsmith RC, Riggs AC, D’Antonio CM. 1980. Survival of hurricane-generated coral fragments and a disturbance model of reef calcification/growth rates. Oecologia 46:322–329. doi:10.1007/BF00346259

Hijmans RJ, Phillips S, Leathwick JR, Elith J. 2017. dismo: species distribution modeling. R Packag version 1.1-4.

Hogeweg P, Hesper B. 1990. Crowns crowding: an individual oriented model of the Acanthaster phenomenon In: Bradbury R, editor. Acanthaster and the Coral Reef: A Theoretical Perspective. Lecture Notes in Biomathematics Volume 88. Springer, Berlin, Heidelberg. pp. 169–188. doi:10.1007/978-3-642-46726-4_11

Holbrook SJ, Adam TC, Edmunds PJ, Schmitt RJ, Carpenter RC, Brooks AJ, Lenihan HS, Briggs CJ. 2018. Recruitment Drives Spatial Variation in Recovery Rates of Resilient Coral Reefs. Sci Rep 8:1–11. doi:10.1038/s41598-018-25414-8

Huang D, Roy K. 2015. The future of evolutionary diversity in reef corals. Philos Trans R Soc B 370:20140010. doi:10.1098/rstb.2014.0010

Hughes TP, Graham NAJ, Jackson JB. C, Mumby PJ, Steneck RS. 2010. Rising to the challenge of sustaining coral reef resilience. Trends Ecol Evol 25:633–42. doi:10.1016/j.tree.2010.07.011

Hughes TP, Kerry JT, Baird AH, Connolly SR, Chase TJ, Dietzel A, Hill T, Hoey AS, Hoogenboom MO, Jacobson M, Kerswell A, Madin JS, Mieog A, Paley AS, Pratchett MS, Torda G, Woods RM. 2019. Global warming impairs stock– recruitment dynamics of corals. Nature 568:387–390. doi:10.1038/s41586-019-1081-y

Hughes TP, Kerry JT, Baird AH, Connolly SR, Dietzel A, Eakin CM, Heron SF, Hoey AS, Hoogenboom MO, Liu G, McWilliam MJ, Pears RJ, Pratchett MS, Skirving WJ, Stella JS, Torda G. 2018. Global warming transforms coral reef assemblages. Nature 556:492–496. doi:10.1038/s41586-018-0041-2

Johnson DW, Christie MR, Pusack TJ, Stallings CD, Hixon MA. 2018. Integrating larval connectivity with local demography reveals regional dynamics of a marine metapopulation. Ecology 99:1419–1429. doi:10.1002/ecy.2343

Kayal M, Lenihan HS, Brooks AJ, Holbrook SJ, Schmitt RJ, Kendall BE. 2018. Predicting coral community recovery using multi-species population dynamics models. Ecol Lett 21:1790–1799. doi:10.1111/ele.13153

Kennedy E V, Perry CT, Halloran PR, Iglesias-Prieto R, Scho CHL, Wisshak M, Form AU, Carricart-ganivet JP, Fine M, Eakin CM, Mumby PJ, Schönberg CHL, Wisshak M, Form AU, Carricart-ganivet JP, Fine M, Eakin CM, Mumby PJ. 2013. Avoiding coral reef functional collapse requires local and global action. Curr Biol 23:912–8. doi:10.1016/j.cub.2013.04.020

Kubicek A, Borell E. 2011. Modelling resilience and phase shifts in coral reefs: Application of different modelling approaches In: Jopp F, Reuter H, Breckling B, editors. Modelling Complex Ecological Dynamics. Berlin, Heidelberg: Springer Berlin Heidelberg. pp. 241–255. doi:10.1007/978-3-642-05029-9

Kubicek A, Jopp F, Breckling B, Lange C, Reuter H. 2015. Context-oriented model validation of individual-based models in ecology: A hierarchically structured approach to validate qualitative, compositional and quantitative characteristics. Ecol Complex 22:178–191. doi:10.1016/j.ecocom.2015.03.005

Kubicek A, Muhando C, Reuter H. 2012. Simulations of long-term community dynamics in coral reefs - How perturbations shape trajectories. PLoS Comput Biol 8:e1002791. doi:10.1371/journal.pcbi.1002791

Kubicek A, Reuter H. 2016. Mechanics of multiple feedbacks in benthic coral reef communities. Ecol Modell 329:29–40. doi:10.1016/j.ecolmodel.2016.02.018

Langmead O, Sheppard C. 2004. Coral reef community dynamics and disturbance: a simulation model. Ecol Modell 175:271–290. doi:10.1016/j.ecolmodel.2003.10.019

Loreau M. 2000. Biodiversity and ecosystem functioning: recent theoretical advances. Okios 91:3–17. doi:10.1034/j.1600-0706.2000.910101.x

Madin JS, Anderson KD, Andreasen MH, Bridge TCL, Cairns SD, Connolly SR, Darling ES, Diaz M, Falster DS, Franklin EC, Gates RD, Hoogenboom MO, Huang D, Keith SA, Kosnik MA, Kuo C-Y, Lough JM, Lovelock CE, Luiz O, Martinelli J, Mizerek T, Pandolfi JM, Pochon X, Pratchett MS, Putnam HM, Roberts TE, Stat M, Wallace CC, Widman E, Baird AH. 2016a. The Coral Trait Database, a curated database of trait information for coral species from the global oceans. Sci Data 3:160017. doi:10.1038/sdata.2016.17

Madin JS, Connolly SR. 2006. Ecological consequences of major hydrodynamic disturbances on coral reefs. Nature 444:477–80. doi:10.1038/nature05328

Madin JS, Dell AI, Madin EMP, Nash MC. 2012a. Spatial variation in mechanical properties of coral reef substrate and implications for coral colony integrity. Coral Reefs 32:173–179. doi:10.1007/s00338-012-0958-0

Madin JS, Hoogenboom MO, Connolly SR, Darling ES, Falster DS, Huang D, Keith SA, Mizerek T, Pandolfi JM, Putnam HM, Baird AH. 2016b. A trait-based approach to advance coral reef science. Trends Ecol Evol 31:419–428. doi:10.1016/j.tree.2016.02.012

Madin JS, Hughes TP, Connolly SR. 2012b. Calcification, storm damage and population resilience of tabular corals under climate change. PLoS One 7:e46637. doi:10.1371/journal.pone.0046637

McCann KS. 2000. The diversity-stability debate. Nature 405:228–33. doi:10.1038/35012234

Mcleod E, Anthony KRN, Mumby PJ, Maynard J, Beeden R, Graham NAJJ, Heron SF, Hoegh-Guldberg O, Jupiter S, Macgowan P, Mangubhai S, Marshall N, Marshall PA, McClanahan TR, Mcleod K, Nyström M, Obura D, Parker B, Possingham HP, Salm R V., Tamelander J. 2019. The future of resilience-based management in coral reef ecosystems, Journal of Environmental Management. Academic Press. doi:10.1016/j.jenvman.2018.11.034

McWilliam M, Chase TJ, Hoogenboom MO. 2018a. Neighbor diversity regulates the productivity of coral assemblages. Curr Biol 28:3634–3639.e3. doi:10.1016/j.cub.2018.09.025

McWilliam M, Hoogenboom MO, Baird AH, Kuo C, Madin JS, Hughes TP. 2018b. Biogeographical disparity in the functional diversity and redundancy of corals. Proc Natl Acad Sci 1–6. doi:10.1073/pnas.1716643115

Melbourne-Thomas J, Johnson CR, Fulton EA. 2011a. Regional-scale scenario analysis for the Meso-American Reef system: Modelling coral reef futures under multiple stressors. Ecol Modell 222:1756–1770. doi:10.1016/j.ecolmodel.2011.03.008

Melbourne-Thomas J, Johnson CR, Fung T, Seymour RM, Chérubin LM, Arias-González JE, Fulton EA, Seymour RM, Chérubin LM, Arias-González JE, Fulton EA. 2011b. Regional-scale scenario modeling for coral reefs: A decision support tool to inform management of a complex system. Ecol Appl 21:1380–1398. doi:10.1890/09-1564.1

Mellin C, Bradshaw CJA, Fordham DA, Caley MJ. 2014. Strong but opposing β-diversity– stability relationships in coral reef fish communities. Proc R Soc B 281:20131993. doi:10.1098/rspb.2013.1993

Moberg F, Folke C. 1999. Ecological goods and services of coral reef ecosystems. Ecol Econ 29:215–233. doi:10.1016/S0921-8009(99)00009-9

Mumby PJ. 2006. The impact of exploiting grazers (Scaridae) on the dynamics of Caribbean coral reefs. Ecol Appl 16:747–69. doi:10.1890/1051-0761(2006)016[0747:TIOEGS]2.0.CO;2

Mumby PJ, Hastings A, Edwards HJ. 2007. Thresholds and the resilience of Caribbean coral reefs. Nature 450:1–10. doi:10.1038/nature06252

Mumby PJ, Steneck RS. 2008. Coral reef management and conservation in light of rapidly evolving ecological paradigms. Trends Ecol Evol 23:555–63. doi:10.1016/j.tree.2008.06.011

Muthukrishnan R, Lloyd-Smith JO, Fong P. 2016. Mechanisms of resilience: empirically quantified positive feedbacks produce alternate stable states dynamics in a model of a tropical reef. J Ecol 104:1662–1672. doi:10.1111/1365-2745.12631

Nash KL, Graham NAJ, Jennings S, Wilson SK, Bellwood DR. 2016. Herbivore cross-scale redundancy supports response diversity and promotes coral reef resilience. J Appl Ecol 53:646–655. doi:10.1111/1365-2664.12430

Nelson HR, Kuempel CD, Altieri AH. 2016. The resilience of reef invertebrate biodiversity to coral mortality. Ecosphere 7:1–14. doi:10.1002/ecs2.1399

Newman SP, Meesters EH, Dryden CS, Williams SM, Sanchez C, Mumby PJ, Polunin NVC. 2015. Reef flattening effects on total richness and species responses in the Caribbean. J Anim Ecol 84:1678–1689. doi:10.1111/1365-2656.12429

North MJ, Collier NT, Ozik J, Tatara ER, Macal CM, Bragen M, Sydelko P. 2013. Complex adaptive systems modeling with Repast Simphony. Complex Adapt Syst Model 26. doi:10.1186/2194-3206-1-3

Ortiz JC, Bozec Y-MM, Wolff NH, Doropoulos C, Mumby PJ. 2014. Global disparity in the ecological benefits of reducing carbon emissions for coral reefs. Nat Clim Chang 4:1090–1094. doi:10.1038/nclimate2439

Paradis E, Schliep K. 2019. Ape 5.0: An environment for modern phylogenetics and evolutionary analyses in R. Bioinformatics 35:526–528. doi:10.1093/bioinformatics/bty633

Parrott L. 2002. Complexity and the limits of ecological engineering. Am Soc Agric Eng 45:1–6. doi:10.13031/2013.11032

Perry CT, Murphy GN, Kench PS, Edinger EN, Smithers SG, Steneck RS, Mumby PJ. 2014. Changing dynamics of Caribbean reef carbonate budgets: Emergence of reef bioeroders as critical controls on present and future reef growth potential. Proc R Soc B Biol Sci 281. doi:10.1098/rspb.2014.2018

Perry CT, Murphy GN, Kench PS, Smithers SG, Edinger EN, Steneck RS, Mumby PJ. 2013. Caribbean-wide decline in carbonate production threatens coral reef growth. Nat Commun 4:1–7. doi:10.1038/ncomms2409

Pratchett MS, Hoey AS, Wilson SK. 2014. Reef degradation and the loss of critical ecosystem goods and services provided by coral reef fishes. Curr Opin Environ Sustain 7:37–43. doi:10.1016/j.cosust.2013.11.022

Pratchett MS, Thompson CA, Hoey AS, Cowman PF, Wilson SK. 2018. Effects of coral bleaching and coral loss on the structure and function of reef fish assemblages In: van Oppen M, Lough J, editors. Coral Bleaching. Ecological Studies (Analysis and Synthesis), Vol 233. Springer, Cham. pp. 265–293. doi:10.1007/978-3-319-75393-5_11

Prowse TAA, Bradshaw CJA, Delean S, Cassey P, Lacy RC, Wells K, Aiello-Lammens ME, Akçakaya HR, Brook BW. 2016. An efficient protocol for the global sensitivity analysis of stochastic ecological models. Ecosphere 7:1–17. doi:10.1002/ecs2.1238

R-Core Team. 2017. R: A language and environment for statistical computing.

Revell LJ. 2012. phytools: an R package for phylogenetic comparative biology (and other things). Methods Ecol Evol 3:217–223. doi:10.1111/j.2041-210X.2011.00169.x

Roff G, Doropoulos C, Zupan M, Rogers A, Steneck RS, Golbuu Y, Mumby PJ. 2015. Phase shift facilitation following cyclone disturbance on coral reefs. Oecologia. doi:10.1007/s00442-015-3282-x

Salguero-Gómez R, Violle C, Gimenez O, Childs D. 2018. Delivering the promises of trait-based approaches to the needs of demographic approaches, and vice versa. Funct Ecol 32:1424–1435. doi:10.1111/1365-2435.13148

Sammarco PW. 1980. Diadema and its relationship to coral spat mortality: grazing, competition, and biological disturbance. J Exp Mar Bio Ecol 45:245–272. doi:10.1016/0022-0981(80)90061-1

Sandin SA, McNamara DE. 2012. Spatial dynamics of benthic competition on coral reefs. Oecologia 168:1079–1090. doi:10.1007/s00442-011-2156-0

Scheffer M, Carpenter S, Foley JA, Folke C, Walker B. 2001. Catastrophic shifts in ecosystems. Nature 413:591–596. doi:10.1038/35098000

Scheffer M, Carpenter SR. 2003. Catastrophic regime shifts in ecosystems: linking theory to observation. Trends Ecol Evol 18:648–656. doi:10.1016/j.tree.2003.09.002

Sheppard C, Dixon DJ, Gourlay M, Sheppard A, Payet R. 2005. Coral mortality increases wave energy reaching shores protected by reef flats: examples from the Seychelles. Estuar Coast Shelf Sci 64:223–234. doi:10.1016/j.ecss.2005.02.016

Sleeman JC, Boggs GS, Radford BC, Kendrick GA. 2005. Using agent-based models to aid reef restoration: enhancing coral cover and topographic complexity through the spatial arrangement of coral transplants. Restor Ecol 13:685–694. doi:10.1111/j.1526-100X.2005.00087.x

Stekhoven DJ, Bühlmann P. 2012. Missforest - nonparametric missing value imputation for mixed-type data. Bioinformatics 28:112–118. doi:10.1093/bioinformatics/btr597

Swain TD, Vega-Perkins JB, Oestreich WK, Triebold C, DuBois E, Henss J, Baird A, Siple M, Backman V, Marcelino L. 2016. Coral bleaching response index: a new tool to standardize and compare susceptibility to thermal bleaching. Glob Chang Biol 22:2475–2488. doi:10.1111/gcb.13276

Tam T, Ang PO. 2012. Object-oriented simulation of coral competition in a coral reef community. Ecol Modell 245:111–120. doi:10.1016/j.ecolmodel.2012.03.023

Tam T, Ang PO. 2009. Catastrophic regime shifts in coral communities exposed to physical disturbances: simulation results from object-oriented 3-dimensional coral reef model. J Theor Biol 259:193–208. doi:10.1016/j.jtbi.2009.03.014

Thibaut LM, Connolly SR, Sweatman HPA. 2012. Diversity and stability of herbivorous fishes on coral reefs. Ecology 93:891–901. doi:10.1890/11-1753.1

Torda G, Sambrook K, Cross P, Sato Y, Bourne DG, Lukoschek V, Hill T, Torras Jorda G, Moya A, Willis BL. 2018. Decadal erosion of coral assemblages by multiple disturbances in the Palm Islands, central Great Barrier Reef. Sci RePoRTS | 8:11885. doi:10.1038/s41598-018-29608-y

Urbanek S. 2018. rJava: low-level R to Java interface. R Packag version 0.9-10.

van de Leemput IA, Hughes TP, van Nes EH, Scheffer M. 2016. Multiple feedbacks and the prevalence of alternate stable states in coral reefs. Coral Reefs 1–22. doi:10.1007/s00338-016-1439-7

van der Laan JD, Bradbury RH, Bradbury VDL. 1990. Futures of the Great Barrier Reef ecosystem. Math Comput Model 14:705–709. doi:10.1016/0895-7177(90)90273-P

Violle C, Navas ML, Vile D, Kazakou E, Fortunel C, Hummel I, Garnier E. 2007. Let the concept of trait be functional! Oikos 116:882–892. doi:10.1111/j.2007.0030-1299.15559.x

Wakeford M, Done TJ, Johnson CR. 2008. Decadal trends in a coral community and evidence of changed disturbance regime. Coral Reefs 27:1–13. doi:10.1007/s00338-007-0284-0

Wear SL, Thurber RV. 2015. Sewage pollution: Mitigation is key for coral reef stewardship. Ann N Y Acad Sci 1355:15–30. doi:10.1111/nyas.12785

Weijerman M, Fulton EA, Janssen ABG, Kuiper JJ, Leemans R, Robson BJ, van de Leemput IA, Mooij WM. 2015. How models can support ecosystem-based management of coral reefs. Prog Oceanogr 138:559–570. doi:10.1016/j.pocean.2014.12.017

Wilkinson C. 2008. Status of Coral Reefs of the World: 2008, Global Coral Reef Monitoring Network and Reef and Rainforest Research Centre. Towsnville, Australia.

Williams I, Polunin N. 2001. Large-scale associations between macroalgal cover and grazer biomass on mid-depth reefs in the Caribbean. Coral Reefs 19:358–366. doi:10.1007/s003380000121

Wolanski E, Kingsford M. 2014. Oceanographic and behavioural assumptions in models of coral reef larval fish oceanography. J R Soc Interface 11:1–19. doi:10.1098/rsif.2014.0209

Wolanski E, Richmond RH, McCook L. 2004. A model of the effects of land-based, human activities on the health of coral reefs in the Great Barrier Reef and in Fouha Bay, Guam, Micronesia. J Mar Syst 46:133–144. doi:10.1016/j.jmarsys.2003.11.018

Woodhead AJ, Hicks CC, Norström A V., Williams GJ, Graham NAJ. 2019. Coral reef ecosystem services in the Anthropocene. Funct Ecol 33:1023–1034. doi:10.1111/1365-2435.13331

Zaneveld JR, Burkepile DE, Shantz AA, Pritchard CE, McMinds R, Payet JP, Welsh R, Correa AMS, Lemoine NP, Rosales S, Fuchs C, Maynard JA, Vega Thurber RL. 2016. Overfishing and nutrient pollution interact with temperature to disrupt coral reefs down to microbial scales. Nat Commun 1–12. doi:10.1038/ncomms11833

Zeileis A, Hothorn T. 2002. Diagnostic checking in regression relationships. R News 2:7–10.

Zhang SY, Speare KE, Long ZT, McKeever KA, Gyoerkoe M, Ramus AP, Mohorn Z, Akins KL, Hambridge SM, Graham NAJ, Nash KL, Selig ER, Bruno JF. 2014. Is coral richness related to community resistance to and recovery from disturbance? PeerJ 2:e308. doi:10.7717/peerj.308

